## Supplemental Figures for "Chromosome-specific differences in the recombination landscape of spontaneous meiotic nondisjunction"

### Supplementary Information

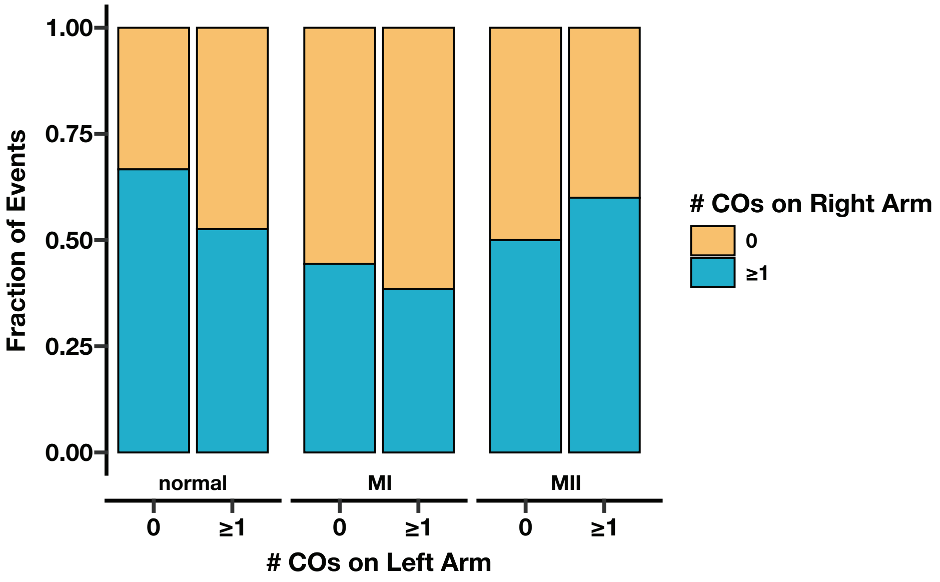


**Figure S1. The influence of presence of crossovers on the left arm of chromosome *2* on the presence of crossovers on the right arm appears minimally affected by meiotic NDJ.**
The fraction of normal, MI, and MII meiotic NDJ events with zero or at least one detectable crossover on the left arm of chromosome *2* with zero (orange) or at least one (blue) detectable crossover on the right arm of chromosome *2*. Normal meioses were taken from Miller et al. (2016).

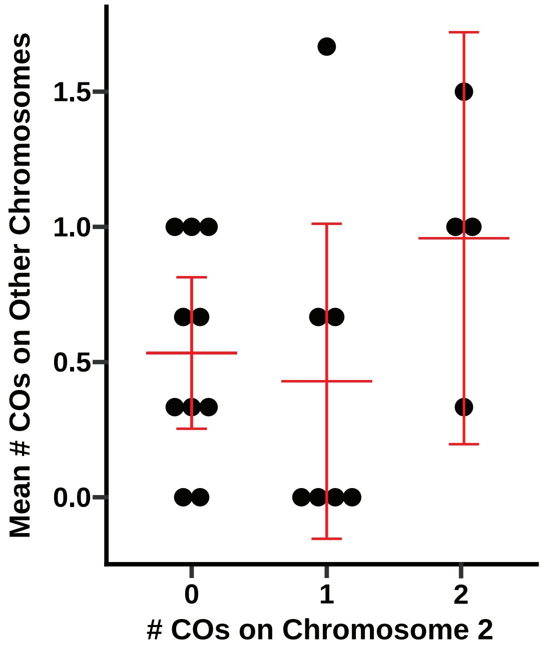


**Figure S2. Number of crossovers on chromosome *2* is minimally correlated with recombination rate on other chromosomes in MI NDJ.** The mean number of crossovers on chromosomes *X* and *3* relative to how many crossovers were detected on chromosome *2* in each MI NDJ event. Large red bars indicate means, with smaller bars indicating 95% confidence intervals.

### Supplementary Tables

**Table S1. *Drosophila* stocks.**
A table of all stocks used. All stocks are from Bloomington Stock Center, and stock numbers for each genotype are provided.

**Table S2. NDJ progeny used for crossover analysis.**
A table of all NDJ progeny with their sequencing run information and barcodes used for sample pooling. Barcode numbers and sequences are consistent with those from Oxford Nanopore Technologies Native Barcoding Kit SQL-NBD114.24.

**Table S3. Crossovers identified in meiotic NDJ progeny.**
All crossovers identified in NDJ progeny are provided with the NDJ male of origin, type of crossover (single or double), and positions of 5' and 3' SNPs used to locate them. In addition, chromosome arms without crossovers are listed with the 5' and 3' SNPs available for that chromosome arm.
